# Time-oriented attention improves accuracy in a paced finger tapping task

**DOI:** 10.1101/2021.04.16.440197

**Authors:** Leonardo Versaci, Rodrigo Laje

**Affiliations:** Departamento de Ciencia y Tecnología, Universidad Nacional de Quilmes, Bernal, Buenos Aires, B1876BXD, Argentina; CONICET, Buenos Aires, C1425FQB, Argentina

**Author notes:** Electronic mail: Corresponding author.

**Keywords:** Sensorimotor synchronization, Finger tapping, Attention, Time processing, Event-related potentials

## Abstract

Finger tapping is a task widely used in a variety of experimental paradigms, in particular to understand sensorimotor synchronization and time processing in the range of hundreds of milliseconds (millisecond timing). Normally, subjects don’t receive any instruction about what to attend to and the results are seldom interpreted taking into account the possible effects of attention. In this work we show that attention can be oriented to the purely temporal aspects of a paced finger tapping task and that it affects performance. Specifically, time-oriented attention improves the accuracy in paced finger tapping and it also increases the resynchronization efficiency after a period perturbation. We use two markers of the attention level: auditory ERPs and subjective report of the mental workload. In addition, we propose a novel algorithm to separate the auditory, stimulus-related components from the somatosensory, response-related ones, which are naturally overlapped in the recorded EEG.

## I. INTRODUCTION

In this work we address the question of whether sensorimotor synchronization is affected by the level of attention during a paced finger tapping task. In a practical way, if the subject’s performance is indeed modified by attention then future experiments should consider attention as a variable to be controlled for, or at least take into account its contribution to the overall variability. In the same sense, a result like that might lead us to reconsider or reinterpret many published results in the literature of paced finger tapping given the ample variety of experimental conditions where attention would be expected to vary. In a more fundamental sense, establishing the role of attention in paced finger tapping and thus deciding whether the task is strongly automatic Kahneman and Chajczyk (1983) might help us identify which brain regions are involved, a still unanswered question due to the multitude of hypothesized processes underlying this behavior López and Laje (2019).

### A. Paced finger tapping and the error correction mechanism

In a paced finger tapping task the subject is instructed to tap in synchrony with a periodic external stimulus, as in keeping pace with music. This very simple task allows us to study the processing of time in the brain Bavassi *et al*. (2017), multisensory integration Mates and Aschersleben (2000), interpersonal coordination Konvalinka *et al*. (2010), timing mechanisms in nonhuman primates Merchant, Harrington, and Meck (2013) and interspecific comparisons Zarco *et al*. (2009), differences between musicians and nonmusicians Franěk *et al*. (1991), metrical perception Iversen *et al*. (2015), motor control Dione and Delevoye-Turrell (2015), and higher cognitive functions like attention Miyake, Onishi, and Pöppel (2004). It is important to note that paced finger tapping cannot be sustained without an error correction mechanism—otherwise the intrinsic variability of motor responses and the smallest detuning between actual and perceived interstimulus interval would accumulate leading to diverging differences between stimuli and taps Repp (2005).

One of the most commonly used observables to study the error correction mechanism is the asynchrony, i.e. the difference between the occurrence time of the *n*-th response *R*_*n*_ (tap) and the occurrence time of the associated stimulus *S*_*n*_ (beep): *e*_*n*_ = *R*_*n*_ – *S*_*n*_. In a paced finger tapping task the mean asynchrony (MA) is usually negative, meaning that on average the response precedes the stimulus. Normally, non musician subjects have a MA of a few tens of milliseconds depending on the experimental condition Mates (1994). For instance, Aschersleben and Prinz Aschersleben and Prinz (1995) observed that additional auditory feedback from the taps produces a decrease in absolute value of the MA (i.e. it gets closer to zero). It is also observed that MA depends on the interstimulus interval (ISI), with MA values getting more negative (i.e. larger in absolute value) as the ISI is increased Kolers and Brewster (1985); Repp (2003). The fact that in general taps precede beeps is one of the most successfully replicated findings in sensorimotor synchronization (SMS), and it is called “negative MA” (NMA) Repp (2005). Surprisingly, there is no clear consensus yet about its mechanisms and the absence of an informed theoretical framework makes its interpretation difficult. On the other hand, the MA has shown to be sensitive to experimental manipulation, which allows us to investigate the factors the error correction mechanism depends on Repp (2005).

### B. The error correction mechanism and the role of perturbations

The error correction mechanism can be probed by performing perturbations to the stimuli sequence. One of the most frequently used temporal perturbations is an abrupt change in the interstimulus interval, known as a step-change (an abrupt change of tempo in musical terms). When unexpected, the temporal perturbation induces a forced error in the asynchrony at the perturbation beep, after which the subject has to recover average synchronization, usually achieved in a few taps. There can be differences between the pre- and post-perturbation steady states. Bavassi *et al*. Bavassi, Tagliazucchi, and Laje (2013) reported a change between pre- and post-perturbation MA opposite to the one expected by a parametric change in ISI of the same size (i.e. between constant ISI conditions). Praamstra and coworkers Praamstra *et al*. (2003) reported a similar effect with phase-shift perturbations.

Between the pre- and post-perturbation steady states there is a resynchronization phase. Resynchronization is the transient that begins with the forced error in the perturbation beep and ends when the subject reaches the post-perturbation baseline. Several measures have been proposed to quantify the resynchronization phase, for instance, the phase correction response (PCR) defined as the difference between the occurrence time of the first response after perturbation and its expected occurrence time if no perturbation was in place Repp and Keller (2004); Repp (2011). In this work we introduce a new measure that takes into account the pre-post change in MA called *resynchronization efficiency*.

### C. Temporal perturbations and temporal attention

The effect of attention on a finger tapping task has been studied under a few conditions. Repp *et al*. Repp and Keller (2004) analyzed the effect of attention on the resynchronization phase after step-change perturbations by using a dual-task paradigm, with arithmetic operations as secondary tasks. He observed that subjects corrected more quickly (greater PCR) in the single-versus dual-task condition. In other words, they concluded that diverting attention from the main (finger tapping) task makes the resynchronization phase slower (smaller PCR) after a step change.

The effect of attention on the steady-state phase in a finger tapping task was analyzed by Miyake *et al*. Miyake, Onishi, and Pöppel (2004) in a dual-task paradigm with word recall as secondary task. He didn’t find any difference in asynchrony variability between single- and dual-task conditions across a range of ISI from 450 ms to 1500 ms. Regarding the steady state phase, Miyake concludes that finger tapping synchronization is an automatic behavior not mediated by attention. A different work by Caspi Caspi (2002) attempted to modify the MA in the steady state by instructing the subject to focus the attention on the tactile sensation from the tap. In this work, Caspi made the tapping surface shift downwards at some point to prevent the finger from making contact and instructed the subjects to stop tapping immediately after perceiving it. His hypothesis was that this instruction would orient attention towards the tactile sensation, probably making it have better temporal resolution and thus decreasing the time difference between tap and beep. Difficulties with the experimental design hampered reaching a clear conclusion.

It is worth noting the purely temporal nature of the paced finger tapping task. It is reasonable thus to expect effects on the performance when the attention is focused specifically on temporal aspects of the task. However, to the best of our knowledge there are no published works on the effects of time-oriented attention on paced finger tapping. In this work we developed a novel experimental paradigm according to this idea.

### D. Attention and electrophysiology

Our experimental paradigm allows us to manipulate the level of attention, and we record auditory ERPs in order to have an electrophysiological support for it. It is known that attention modulates early components of stimulus-induced ERPs Teder *et al*. (1993); Alho *et al*. (1994a); Hansen and Hillyard (1980); Eimer and Forster (2003a). In a paced finger tapping task, however, it is difficult to isolate stimulus-induced ERPs because of contamination with sensorimotor activity from the taps. Due to the very nature of the task, one could expect temporal overlapping of both tap-related components (finger movement, and proprioceptive and tactile perception) and stimulus-related components (auditory stimuli). To address this difficulty we turn to a work by Woldorff Woldorff (1993) who developed the Adjar algorithm to isolate adjacent ERPs. The algorithm was originally designed for an experimental paradigm with dichotic auditory stimuli of very short ISIs, but it is powerful enough to be generalized. In this work we developed a novel algorithm to isolate the auditory stimulus-induced ERPs in a paced finger tapping task by adapting Woldorff’s algorithm (see details in the Appendix).

### E. This work

We set out to analyze the synchronization performance in a paced finger tapping task with perturbations during both the steady-state and resynchronization phases to study the potential effects of time-oriented attention. To this we set two levels of attention: NORMAL and HIGH. Unlike other works Caspi (2002); Miyake, Onishi, and Pöppel (2004); Repp and Keller (2004), our HIGH condition makes the subject direct the attention to the temporal aspects of the stimuli sequence. We test the hypothesis of whether attention improves performance in two potential ways: that a higher level of attention leads to better accuracy and precision in the steady state phase, and to a higher efficiency in the resynchronization phase. In addition, two more factors are included in the experimental design: auditory feedback and perturbation sign. Finally, to have a quantitative support for our claim of an actually higher level of attention in the HIGH condition, we report two measures commonly used as evidence of a change in attention level: auditory ERPs and mental workload.

## II. METHODS

### A. Subjects and ethical considerations

Forty-four right-handed musicians (11 women) participated as paid volunteers ($5, roughly 150 Argentine pesos) after signing an informed consent. We decided to study musicians only (with at least 3 years of practice) because they usually show smaller variability Repp (2010). Mean age was 27.5 years (range 19-44) and mean practice was 5.3 years (range 3-10). We recorded behavioral data (response occurrence times) from all 44 subjects; in addition, we recorded EEG and mental workload from 22 of them. Our experimental protocols were designed in accordance with national and international guidelines and were approved by the Ethics Committee of the University of Quilmes.

### B. Experimental design

We designed a full-factorial experiment, with factors Attention (levels NORMAL and HIGH, between-subjects), Auditory Feedback (levels Without (W/O f) and With (W f), within-subjects), and Perturbation Sign (levels –60 ms and +60 ms, within-subjects). Regarding the Attention factor, subjects were randomly assigned to either one of the levels. The rationale of this design choice was to avoid carryover effects and was based on a pilot experiment where we found that mixing attention levels made all data be contaminated by the HIGH level.

### C. Procedure

Subjects were instructed to tap with the right index finger on a force resistive sensor (FSR) while keeping pace with the metronome. In order to avoid potential differences arising from the effector, all subjects were instructed to use the wrist joint and keep the forearm at rest on the desk. They were told not to move any other body part during a trial and look at a fixation cross in the center of the screen. Before starting the experiment, subjects adapted to the task during a 10-min long practice phase. We created the two levels of attention (NORMAL and HIGH) by using two separate force sensors located side-to-side, in the following way. In the NORMAL condition subjects tapped on the left sensor (see Figure 1A) without any specific instruction about what to pay attention to. In the HIGH condition subjects started tapping on the left sensor, and they had to switch to the right sensor as soon as they perceived the step change perturbation and keep tapping on the right sensor until the end of the trial.

**FIG. 1.**
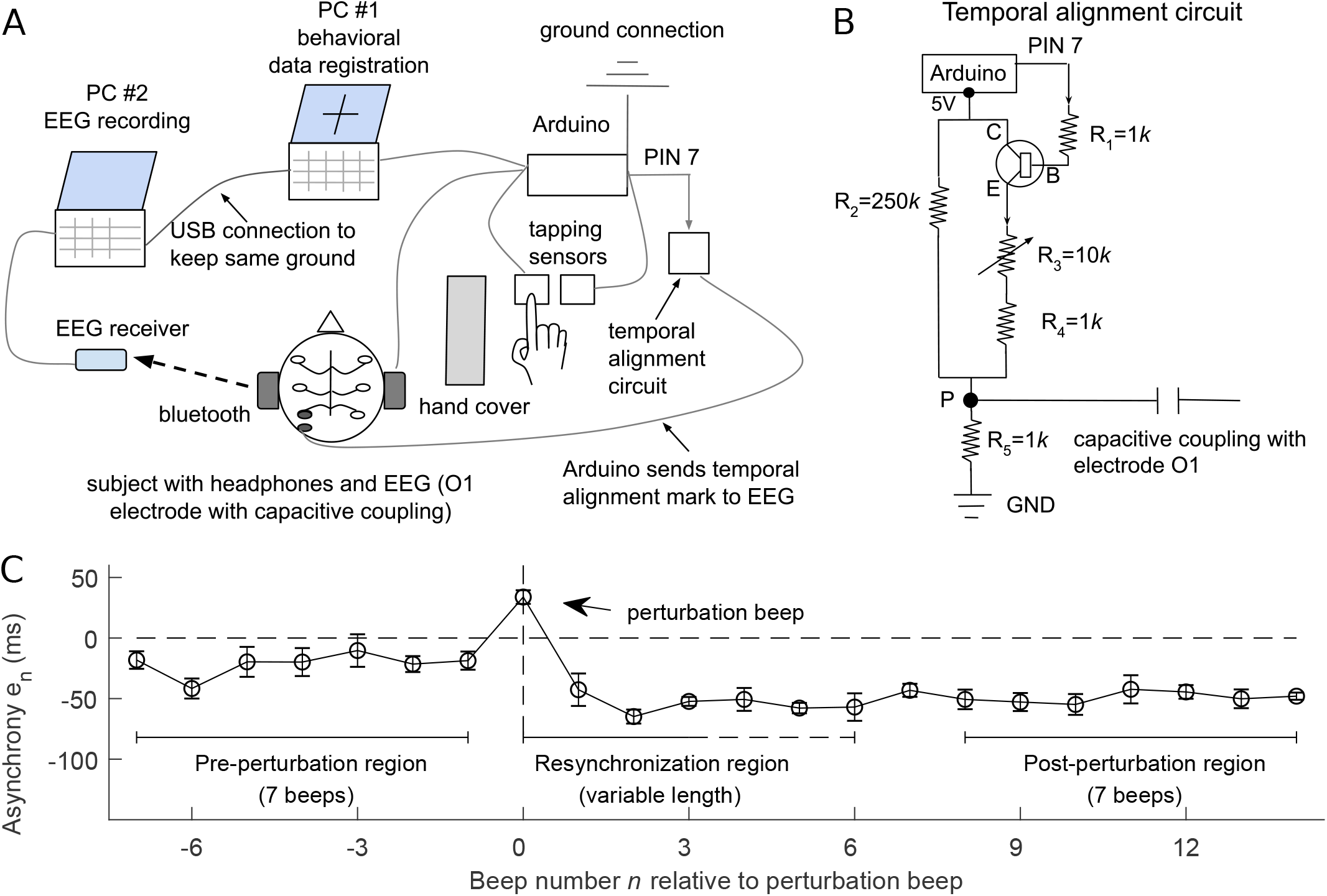
(A) Experimental setup. (B) Circuit for the temporal alignment between Arduino and EEG. (C) Average trial from one subject in one condition (mean *±* standard error across trials).

Every subject was randomly assigned to either NORMAL or HIGH attention. The experiment consisted of two blocks (one block for each feedback condition, in random order and counterbalanced across subjects) with a 3-minute rest between them. Within each block, the subject had to perform 12 valid trials (6 trials from each perturbation sign, random order) with a 1-minute rest after the first 6 valid trials. The experiment’s total duration was approximately 45-60 minutes including the practice phase.

We consider a trial valid if: 1) none of the asynchronies is larger (in absolute value) than 250 ms; 2) there are equal numbers of stimuli and responses since the first recorded response; 3) the first recorded response occurs at most at the third beep; 4) for HIGH attention only: the subject must switch from left sensor to right sensor right after the perturbation (exactly one beep after). If a trial is considered invalid, the subject receives feedback about the type of error and an additional trial is added at the end of the block. The EEG subjects were asked to quantify the mental workload in completing the task at the end of the experiment. According to the Cognitive Load Theory Sweller (2011), the level of automatism in performing a task and associated mental workload are inversely related. We used a 9-point Likert scale, from “very, very low mental effort” (1) through “very, very high mental effort” (9) Paas *et al*. (2003). Each subject reported the mental workload for each feedback condition (W/O f and W f); the mental workload corresponding to the attention level which the subject was assigned to was the average between the two reports.

### D. Behavioral data recording

Stimuli and feedback presentation and response recording were performed with an Arduino Mega 2560 and two force-sensitive resistors (FSR) separated 0.5 cm from each other, connected to PC #1 (the EEG data were recorded by PC #2, see below; see Figure 1A). We used custom-written C code and adapted code from a program originally designed for Arduino Uno Schultz and van Vugt (2016) in order to present sounds with sinusoidal waveform (code). According to Schultz, the combination Arduino-FSR has the shortest latency between the time of contact and the time of occurrence of the feedback (0.6 ms) and also the smallest latency variability (0.3 ms), among other features. Non-EEG subjects used Sennheiser HD 280 Pro headphones; EEG subjects used Sennheiser CX 300-II ear-canal phones. To avoid visual feedback from their own finger movement, the subjects were asked to fix gaze at a cross in the screen and a vertical cardboard screen was put to block peripheral vision.

Each trial was a sequence of 32 stimuli; the pre-perturbation value of the ISI was 600 ms; the perturbation was a tempo step change of ±60 ms, occurring between beeps 11 and 15 at random. Stimulus tones were 50-ms long, 500-Hz sinusoidal waveform sounds; feedback tones were 50-ms long, 1800-Hz sinusoidal waveform sounds. Subjects set sound volume to a comfortable level. The code for controlling the experiment and the code for data analysis were written in MATLAB®.

### E. Behavioral Data Processing

Every subject performed six valid trials for each condition (feedback and perturbation sign). After the experiment we took all valid trials from all subjects and aligned them at the perturbation beep (labeled *n* = 0), and selected the range *n* = –7 through *n* = 14 where all trials from all subjects had responses. We discarded any response outside that range, leaving all trials with the same number of responses (22) and the perturbation at the same position (*n* = 0). For each subject and condition we averaged across the six trials to obtain an average trial in which we defined three regions (Figure 1C):

- Pre-perturbation region (the seven responses before perturbation: *n* = –7 through *n* = –1);
- Post-perturbation region (the last seven responses: *n* = 8 through *n* = 14);
- Resynchronization region (variable length: *n* = 0 through the resynchronization beep which is the first of two consecutive beeps whose asynchronies are statistically indistinguishable from the post-perturbation baseline that is the mean asynchrony of all trials in the post-perturbation region).

As described above, we recorded two groups of subjects:

- Year 2018 (22 subjects; behavior only; headphones);
- Year 2019 (22 subjects; behavior + EEG + mental workload; ear-canal phones).

The *pre-perturbation mean asynchrony* (*MA*_*pre*_) was computed as follows: for each subject and condition, we first averaged the asynchronies in the pre-perturbation region (along the trial), then averaged across trials. Analogously, we defined the *post-perturbation mean asynchrony* (*MA*_*post*_) in the post-perturbation region. We defined the pre-post change in MA as ∆*MA* = *MA*_*post*_ – *MA*_*pre*_.

The *pre-perturbation intratrial SD* (*SD* _*pre*_) was computed as follows: for each subject and condition, we first computed the standard deviation of asynchronies in the pre-perturbation region (along the trial), then averaged across trials. The *post-perturbation intratrial SD* (*SD* _*post*_) follows an analogous definition in the post-perturbation region. We defined the pre-post change in SD as ∆*SD* = *SD* _*post*_ – *SD* _*pre*_.

#### Data pooling

The experimental design is the same for both subject groups (full factorial experiment, three-factor design: attention, feedback, and perturbation sign). To determine whether the groups were comparable we pooled all subjects and performed two ANOVAs, one for *MA*_*pre*_ and one for *SD* _*pre*_ with Year as an additional factor (which represents all differences: year, measurement, and headphone/earphone; we dropped the factor perturbation sign as the two tested values were pre-perturbation only). We didn’t find any significant main or interaction effects of year, either for *MA*_*pre*_ (three-way ANOVA, with year and attention as between-subject factors, and feedback as within-subject factor; significant effect of attention *F* (1, 40) = 11.99, *p* = 0.0013; significant effect of feedback *F* (1, 40) = 5.91, *p* = 0.019; non significant effect of year *F* (1, 40) = 0.05, *p* = 0.81; non significant att x fbk interaction *F* (1, 40) = 0.51, *p* = 0.47; non significant att x year interaction *F* (1, 40) = 0.27, *p* = 0.60; non significant fbk x year interaction *F* (1, 40) = 0.81, *p* = 0.37; non significant att x fbk x year interaction *F* (1, 40) = 0.51, *p* = 0.48) or *SD pre* (three-way ANOVA, with att and year as between-subject factors, and feedback as within-subject factor; non significant effect of attention *F* (1, 40) = 3.53, *p* = 0.06; non-significant effect of feedback *F* (1, 40) = 1.39, *p* = 0.24; non-significant effect of year *F* (1, 40) = 1.72, *p* = 0.19; non-significant att x fbk interaction *F* (1, 40) = 0.02, *p* = 0.87;

non-significant att x year interaction *F* (1, 40) = 0.01, *p* = 0.91; non-significant fbk x year interaction *F* (1, 40) = 2.82, *p* = 0.10; non-significant att x fbk x year interaction *F* (1, 40) = 0.40, *p* = 0.52). Based on these results, we decided to pool all subjects into a single group of *N* = 44.

#### Heteroscedasticity in *MA*_*pre*_

A Levene test applied on MA in the pre-perturbation region revealed a significant heteroscedasticity: inter-subject variability in NORMAL was larger than in HIGH (*p* = 0.019; feedback conditions were averaged). This fact can be observed in Figure 2 where temporal series in HIGH are distributed in a narrower band of asynchronies than in NORMAL. To remove heteroscedasticity a log_10_(–*MA* + constant) transformation was applied. All subsequent analyses and statistical tests were performed on the transformed data.

**FIG. 2.**
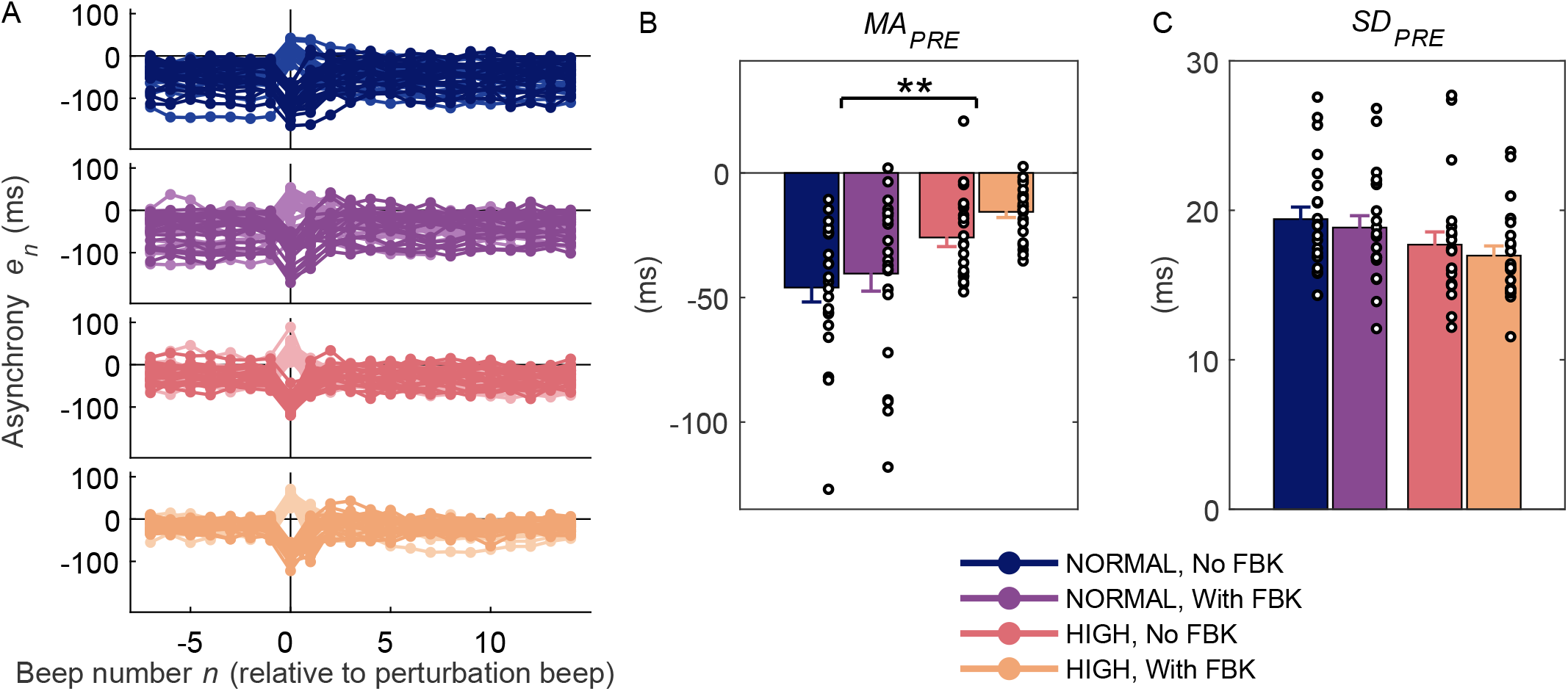
(A) Average time series for each subject and each condition (mean across trials). (B) Attention significantly improves accuracy. Auditory feedback shows a similar behavior but its effect is not significant. Mean asynchrony in the pre-perturbation region *MA*_*pre*_ (mean *±* standard error across subjects). \*\**p* = 0.003. (C) Attention and feedback seem to improve precision too but their effects are not significant. Intra-trial standard deviation in the pre-perturbation region *SD*_*pre*_ (mean *±* standard error across subjects).

#### Expected ∆*MA* and ∆*SD* values from previous behavioral works

The hatched bars in Figure 4 represent the expected values based on a work by Repp Repp (2003) (Figure 2, panels A and B, “Slow auditory” condition) in which the ISI was parametrically varied. We digitized the data and performed a linear regression over an ISI range. The obtained slope for MA was –0.039; therefore the expected change in MA due to an ISI step change of ±60 ms is ±2.34 ms (Figure 4A). The obtained slope for SD was 0.026; thus the expected change in SD due to an ISI step change of ±60 ms is ±1.56 ms (Figure 4B).

### F. Resynchronization efficiency

We define resynchronization efficiency *γ* as:

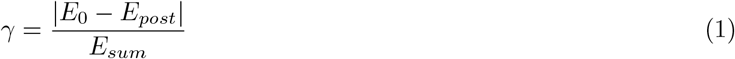

where *E*_0_ is the asynchrony at the perturbation beep, averaged across trials (i.e. the mean forced error at *n* = 0); *E*_*post*_ is the post-perturbation mean asynchrony, averaged across trials; and *E*_*sum*_ is the sum of all asynchrony values (relative to *E*_*post*_) from the perturbation beep (*n* = 0) through the resynchronization beep (*n*_*R*_):

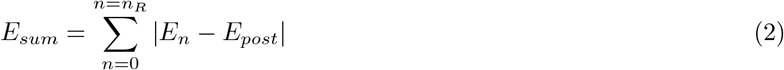

where *E*_*n*_ is the mean asynchrony of the beep *n* across trials in the resynchronization region.

Conceptually, the resynchronization efficiency is a ratio between the time the subject must correct during resynchronization and the sum of actual asynchronies until resynchronized. The resynchronization efficiency takes values ranging from 0 to 1: at fixed value of the forced error *E*_0_ – *E*_*post*_, the shorter the resynchronization phase the greater the efficiency; while at fixed duration of the resynchronization phase, the larger the forced error *E*_0_ – *E*_*post*_ the greater the efficiency. Efficiency also depends on the shape of the asynchrony time series: At fixed forced error and duration of the resynchronization phase, a subject with a shallower asynchrony time series will have a greater efficiency than a subject with constantly high asynchrony values during resynchronization.

### G. EEG recording

EEG recording was performed with an Emotiv EPOC+ system with a sampling rate of 128 Hz. The system has 14 channels placed in locations consistent with the 10-20 montage, and two other additional electrodes located in the mastoids. The Emotiv EPOC+ was specifically validated for auditory ERP recording Badcock *et al*. (2013, 2015) and also for a wide range of measurements Pietto *et al*. (2018). In order to refer the EEG data and the behavioral data to a common time axis, we adapted the Emotiv EPOC+ to receive temporal alignment marks. These marks were sent at the beginning of each trial to one electrode exclusively used to that purpose. The O1 electrode was chosen to accomplish this function because it is located outside the area of study. The 13 remaining electrodes were used to collect the EEG data.

The temporal alignment marks were delivered with a simple, custom-made circuit (Figure 1B). Each mark consists of a sudden variation of the voltage at node P (placed near O1) and is registered by the Emotiv as a well defined oscillation in the O1 channel by means of capacitive coupling. The amplitude of the mark registered in the O1 channel is proportional to the variation of voltage in node P. The circuit consists of a voltage divider in which one of the resistors varies its value according to the state of the kn2222 transistor. When the transistor is in cutoff mode the whole current flows through R2, and the voltage in point P reaches its minimum. When the transistor is in saturation mode most of the current passes through R3 and R4 (since the sum of both resistors is much lower than R2) and the voltage in node P reaches its maximum. The resistor values were selected in such a way that the voltage in node P has minimum and maximum values of 20 mV and 1.5 V, respectively. R3 is a variable resistor used to control the maximum voltage in point P (mark amplitude). The Arduino was programmed to alternate the transistor mode between cutoff (PIN 7=LOW) and saturation (PIN 7=HIGH) every 100 ms during 400 ms. This specific mark is detected by software in a later stage (i.e. offline, after the experiment).

When the finger makes contact with the sensor an electrostatic charge is generated which is observed as a tiny voltage variation in the O1 electrode. To eliminate this source of noise it is necessary to equip the circuit with an adequate ground connection and also have the subjects rest their forearms on the desk surface.

### H. EEG signal processing

The collected EEG data was band-pass filtered with a non-causal, 8th-order Butterworth filter with half-power frequencies of 0.6 Hz and 30 Hz. Response-locked epochs (–180 to 710 ms from each response occurrence) were obtained for every response between the first response and the *n* = –1 response, for every trial. After that, a baseline correction was performed on each epoch by subtracting its mean value. We used three criteria for discarding outlier data. First, epochs were discarded when a voltage sample was greater than ±100 *µ*V (after baseline correction). Second, epochs containing eyes blinks and eyes movements artifacts were discarded by implementing the Step Function algorithm Luck (2014) with a 200-ms moving window in steps of 7 ms. In each window, the mean value of the first 100 ms was compared with the mean value of the last 100 ms; when the difference between both values was greater than ±35 *µ*V the whole epoch was discarded. Third, an epoch was discarded if any of its behavioral asynchronies fell outside the range –170 to +80 ms (values greatly exceeding the 1.5 IQR rule). In addition, a whole electrode was discarded if the number of its valid epochs was lower than 50% of its total number of epochs. This criterion was applied to all electrodes placed within the area of interest (AF3, F3, F7, FC5, FC6, F8, F4, AF4). According to this criterion, three subjects presented one invalid electrode; none of them presented more than one invalid electrode per condition. After the discarding process, there remained an average of 107 valid epochs per subject, condition and electrode (equivalent to an average of discarded epochs of 10.3% per subject, condition, and electrode).

Stimulus-locked epochs were obtained from the valid resp-locked epochs (–100 ms to 350 ms from stimulus occurrence) in the pre-perturbation region (*n* = –7 through *n* = –1). On average, 73 stim-locked epochs were obtained per subject, condition, and electrode. Then, an adaptation of the Adjar algorithm (see appendix) was applied over the stim-locked epochs to remove the ERP components not related to the auditory stimuli.

### I. Permutation testing for ERPs

We implemented a whole-subject permutation approach Luck (2014) to test the difference between NORMAL and HIGH ERPs.

We first defined a temporal window in which the ERPs were compared. Previous works consistently suggest a specific window location and also point to a directional difference between the attended vs non-attended ERPs Teder *et al*. (1993); Hansen and Hillyard (1980); Pereira *et al*. (2014); Näätänen, Gaillard, and Mäntysalo (1978). In a selective attention paradigm with auditory dichotic stimuli, Teder finds that in latencies going from N1 to P2 the stimuli-attended ERP is negatively displaced with respect to the ERP of the non-attended stimuli, with the highest difference occurring around 180 ms (for an ISI of 480). Based on this, we defined a window centered in 180 ms and going from 110 to 250 ms. Next, we created a null distribution by random permutation of the data as follows. Eleven subjects were randomly selected from the group of 22 subjects. We computed an average ERP within this subgroup and called it “surrogate NORMAL”. We averaged the ERPs from the remaining 11 subjects and called it “surrogate HIGH”. We performed a non-paired t-test between the “surrogate NORMAL” and “surrogate HIGH” ERPs for every time sample in the entire window of comparison. We found the maximum of all *t* values in the window and called it *t*_*max*_. We repeated this procedure 1000 times to obtain a distribution of 1000 *t*_*max*_. values. Lastly, we performed a non-paired t-test between the actual NORMAL and HIGH ERPs (i.e. without permutation) for every time sample in the comparison window, and labeled the time bin as significant if the actual *t* value was higher than the 95 percentile of the *t*_*max*_ distribution (i.e. one-tailed). The region where we found significant differences is displayed as a shaded rectangle in Figure 5A.

## III. RESULTS

Figure 2A shows the average time series for each subject and each condition. In the pre-perturbation region (*n* = –7 to *n* = –1, both inclusive) the asynchrony is relatively constant, meaning the subject is synchronized. When perturbation occurs (*n* = 0) a forced error is induced and thus the asynchrony changes abruptly. After that, the subject resynchronizes to a new, slightly different baseline in a few taps.

### A. Attention improves accuracy

Consider first the pre-perturbation region. In this region no effect of perturbation sign is expected so we averaged across *±*60 levels. The pre-perturbation mean asynchrony *MA*_*pre*_ shifts towards less negative values (greater accuracy) when either feedback is added or attention is increased (Figure 2B), but only the attention factor is significant. That is, attention improves accuracy, according to what can be expected intuitively. The additional auditory feedback, also in favor to what is expected since it adds information, shows a similar tendency to produce a less negative MA but the effect is not significant (two-way ANOVA, with attention as between-subject factor and feedback as within-subject factor; attention *F* (1, 42) = 9.84, *p* = 0.003; feedback *F* (1, 42) = 1.74, *p* = 0.19; interaction *F* (1, 42) = 0.03, *p* = 0.86). Figure 2C shows the standard deviation of asynchronies in the pre-perturbation region *SD* _*pre*_. Both increasing attention and adding auditory feedback make *SD* _*pre*_ decrease, i.e. both improve precision, although their effects are not significant (two-way ANOVA, with attention as between-subject factor, feedback as within-subject factor, and subject as random factor; attention *F* (1, 42) = 3.56, *p* = 0.061; feedback *F* (1, 42) = 1.35, *p* = 0.250; interaction *F* (1, 42) = 0.02, *p* = 0.879).

In summary, both attention and auditory feedback behave as intuitively expected with regard to accuracy and precision (i.e. both make them improve, albeit some effects are non significant in our study) in the absence of a perturbation.

### B. Attention and feedback have opposite effects in resynchronization

We now show the subject’s behaviour in the resynchronization phase. Perhaps not surprisingly, we found that a high level of attention improves the resynchronization behaviour of the subject, measured as a significant increase of the efficiency to reach the post-perturbation baseline (Figure 3). On the other hand, the resynchronization efficiency diminishes when auditory feedback is added, which is an unexpected result. Furthermore, the resynchronization efficiency is greater after a positive perturbation (increased period) than after a negative one (decreased period), suggesting asymmetric behavior in line with previous reports Bavassi, Tagliazucchi, and Laje (2013); López and Laje (2019) (three-way ANOVA, with attention as between-subject factor and feedback and perturbation sign as within-subject factors; attention *F* (1, 42) = 5.75, *p* = 0.021; feedback *F* (1, 42) = 11.01, *p* = 0.0019; perturbation sign *F* (1, 42) = 6.56, *p* = 0.014; att x fbk interaction *F* (1, 42) = 0.005, *p* = 0.94; att x sign interaction *F* (1, 42) = 0.23, *p* = 0.62; fbk x sign interaction *F* (1, 42) = 1.09, *p* = 0.30; att x fbk x sign interaction *F* (1, 42) = 2.65, *p* = 0.11).

**FIG. 3.**
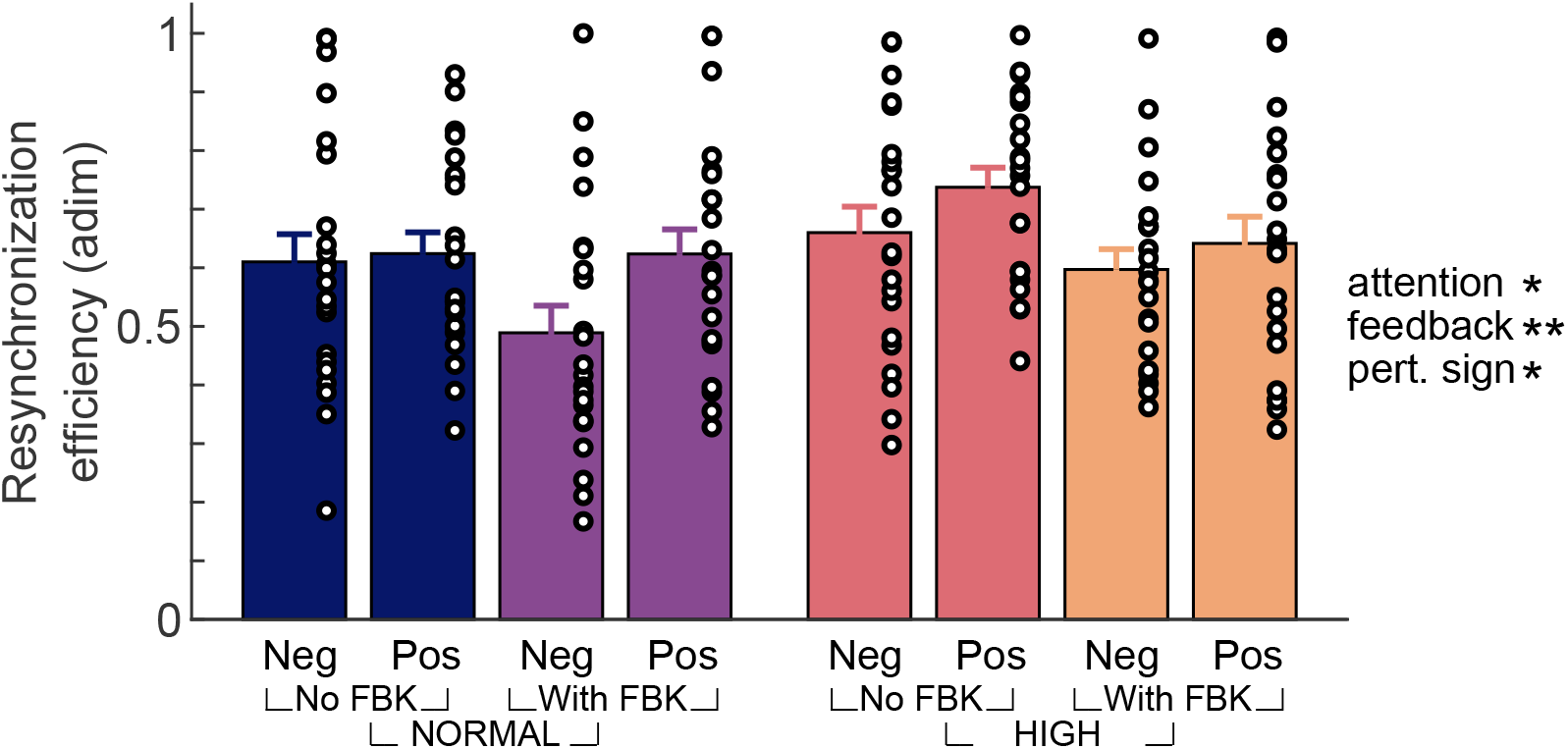
Resynchronization efficiency (mean *±* standard error). Increased attention makes the resynchronization efficiency increase; counterintuitively, the additional feedback makes it decrease. Positive perturbations resynchronize with higher efficiency than negative ones. \**p* = 0.021; \*\**p* = 0.014; \*\*\**p* = 0.0019.

**FIG. 4.**
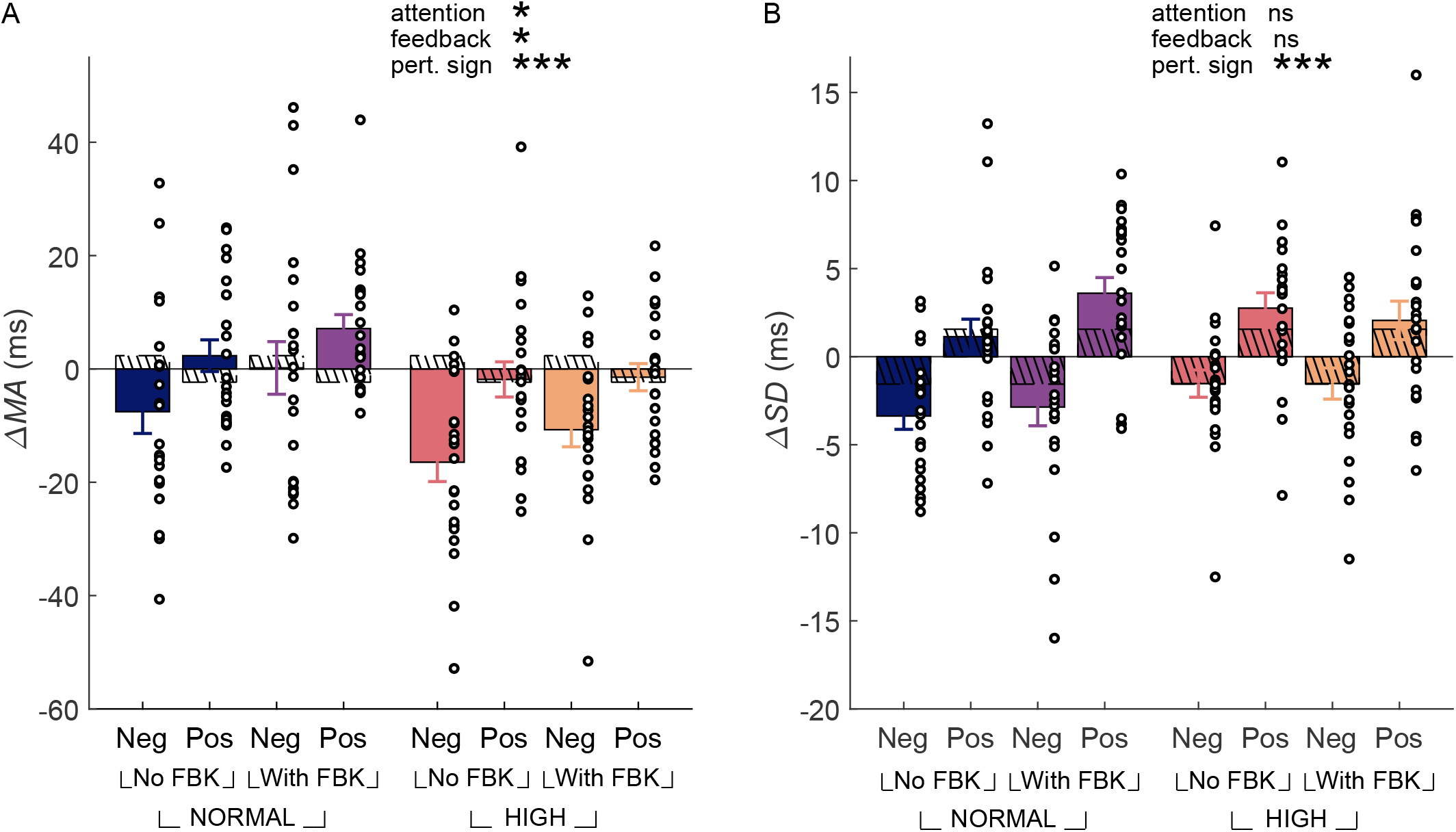
(A) MA varies because of period perturbation, although this variation doesn’t match exactly what we know from works with a parametric manipulation of the period (constant period in each trial). Hatched bars: expected values according to the known dependence of the MA on the period (for example, see Figure 2A in Repp (2003)). \**p* = 0.010; \*\*\**p* = 0.00041. (B) ∆*SD* significantly varies with perturbation sign. After the –60 and +60 perturbations, subjects become more and less precise, respectively. This result roughly agrees with the expected values (hatched bars) from the known dependence of the SD on the period (see Repp (2003), Figure 2B). n/s = non-significant; \*\*\*\**p* = 10^*–*9^.

**FIG. 5.**
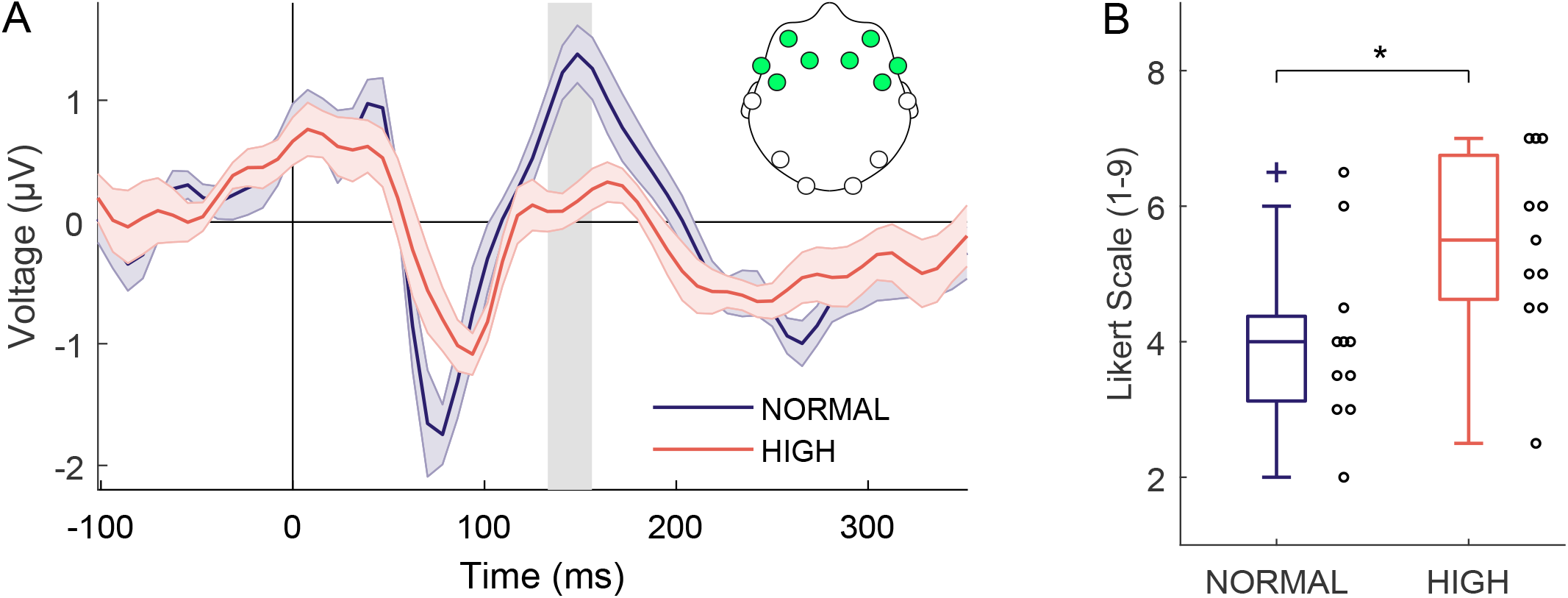
(A) Auditory-related ERPs for NORMAL and HIGH attention in the fronto-central region. The shaded rectangle represents the region where the difference is significant (*t* = 133 ms through 156 ms). Mean *±* standard error across subjects; NORMAL and HIGH levels were averaged across feedback conditions and perturbation signs. (B) Mental workload (subjective report of the mental effort required during the task); \**p* = 0.015.

Notably, none of the interaction effects is significant, which suggests that either adding or removing the auditory feedback doesn’t modify the subject’s attention and allows us to discard a potential confounding (i.e. attention and feedback are independent variables).

### C. Opposite effects of perturbing vs varying the period

We found that a step change perturbation produces a change in the accuracy, in line with our previous reports Bavassi, Tagliazucchi, and Laje (2013) but contrary to what would be expected by parametric manipulation of the period Repp (2003). We first focus on the most common condition in the literature, which corresponds to our NORMAL -W/O f. In that condition, it is known that the MA depends on the period when the period is varied parametrically: isochronous trials with shorter periods make the normally negative MA value get closer to zero (that is, subject more accurate; Figure 2A in Repp (2003)). This contrasts to our previous and current findings: negative perturbations (i.e. where the period is shortened) make the MA shift to more negative values instead (i.e. subject less accurate after perturbation; Figure 4A, first column).

In this work we extend and generalize the observation described above. Figure 4A shows the MA change between the pre- and post-perturbation regions, ∆*MA*, for every experimental condition. All three factors have significant effects on ∆*MA*: on average, high attention leads to a more negative ∆*MA* (i.e. subject less accurate after perturbation), whereas the addition of auditory feedback shifts the post-perturbation MA towards more positive values (subject more accurate after perturbation); the positive perturbations make the MA change less than the negative ones (three-way ANOVA, with attention as between-subject factor and feedback and perturbation sign as within-subject factors; attention *F* (1, 42) = 7.19, *p* = 0.010; feedback *F* (1, 42) = 6.96, *p* = 0.011; perturbation sign *F* (1, 42) = 15.26, *p* = 0.00041; att x fbk interaction *F* (1, 42) = 0.73, *p* = 0.39; att x sign interaction *F* (1, 42) = 0.38, *p* = 0.53; fbk x sign interaction *F* (1, 42) = 1.66, *p* = 0.20; att x fbk x sign interaction *F* (1, 42) = 0.22, *p* = 0.63).

The results in the previous paragraph are not consistent with the values predicted by works where the period was the control parameter (constant in each trial; e.g. Repp (2003), Figure 2A). Those works show that the MA is more negative for larger periods, thus predicting a negative ∆*MA* for perturbations that increase the period (i.e. positive perturbations), and a positive ∆*MA* for perturbations that decrease the period (i.e. negative perturbations as illustrated by hatched bars in Figure 4A). That prediction is almost always opposite to what we get in our experiment. This suggests that a period perturbation has an effect of its own, different from that obtained by parametric manipulation of the period.

Precision is also affected by a step-change perturbation. Figure 4B shows the pre-post change in the intra-trial SD (∆*SD*) for every condition. The perturbation sign is the only parameter with a significant effect: negative perturbations produce a decrease in SD (subjects become more precise after perturbation), while the positive perturbations lead to an increase of SD (subjects less precise after perturbation) (three-way ANOVA, with attention as between-subject factor and feedback and perturbation sign as within-subject factors; attention *F* (1, 42) = 1.43, *p* = 0.23; feedback *F* (1, 42) = 0.80, *p* = 0.37; perturbation sign *F* (1, 42) = 61.1, *p* = 9.9 × 10^*–*10^; att x fbk interaction *F* (1, 42) = 2.08, *p* = 0.15; att x sign interaction *F* (1, 42) = 1.65, *p* = 0.20; fbk x sign interaction *F* (1, 42) = 0.20, *p* = 0.65; att x fbk x sign interaction *F* (1, 42) = 0.92, *p* = 0.34). Parametric manipulation of the period in previous works showed that the SD depends on the period of the trial (Repp (2003), Figure 2B). In our work, only the sign of SD variations was found to be consistent with such dependence; the expected value according to previous reports (hatched bars in Figure 4B) is in general smaller than what we observed. Similarly to what we pointed out for the MA, this suggests that the step-change perturbation might have an effect of its own on the SD.

### D. Electrophysiological correlates of attention level

In order to show that our task makes the subjects reliably change the attention level, we measured two different markers of the level of attention: auditory stimulus-related ERPs (neural correlate of attention), and mental workload (subjective report of mental effort).

Figure 5A shows the auditory stimulus-related ERPs corresponding to each level of attention in the pre-perturbation region (feedback conditions and perturbation signs averaged). As described in the Introduction, in a paced finger tapping task an overlap is expected between auditory (stimulus-related) components and sensorimotor (tap-related) components. According to the literature, the neural correlates of attention rely on the auditory, stimulus-related components so we developed a novel algorithm called TapAdjar (see Appendix) to isolate the auditory components from the whole EEG signal. Significant differences can be observed between attention levels in the P2 component according to what is described in the literature (amplitude of P2 diminishes in high attention conditions; Teder *et al*. (1993); Hansen and Hillyard (1980); Pereira *et al*. (2014); Näätänen, Gaillard, and Mäntysalo (1978); specifically, the HIGH condition shows a large deflection from NORMAL around P2, with significant differences in the shaded rectangle going from 133 to 156 ms after the onset of the stimuli (bootstrapping by *t*_*max*_ permutation approach; see Methods).

The mental workload points in the same direction as shown in Figure 5B: HIGH attention subjects report significantly higher mental effort compared to the NORMAL attention subjects (Likert scale, 1=very very low effort, 9=very very high effort; HIGH: 5.45 ± 1.36, mean±SD; 5.50 ± 1.04, median±MAD. NORMAL: 4.00 ± 1.30, mean±SD; 4 ± 0.90, median±MAD; Kruskal-Wallis test, *p* = 0.015.

## IV. DISCUSSION

### A. SMS is vulnerable to temporal attention

Our results indicate that the error correction mechanism in paced finger tapping may be affected by higher cognitive functions like attention, both in its stationary and resynchronization phases. In the stationary phase attention improves accuracy, while in the resynchronization phase it increases efficiency.

It seems natural to think that higher attention leads to a more accurate behaviour, but it should also be taken into account that attention can be oriented to different aspects of a task, for instance to different sensory modalities or semantic structures stored in memory Posner (1980). The significant effect we found may be related to the fact that attention is oriented specifically to a temporal aspect of the task (tempo change). It is worth considering whether we would observe the same results if attention were oriented to other aspects of the stimuli sequence, like stimulus intensity.

Our results are in clear contrast with Miyake’s hypothesis Miyake, Onishi, and Pöppel (2004) that the error correction mechanism is an automatic process not affected by attention. In fact, our results reveal that the error correction mechanism in a paced finger tapping task is affected by attention when it is oriented to a temporal aspect of the task. It is possible that they didn’t observe any effect of diverting attention because he used a non-temporal secondary task (word recall). Indeed, previous evidence suggests that two temporal tasks performed at once interfere with each other Brown (1997) (although possible interferences between consecutive intervals should be controlled for, see Karmarkar and Buonomano (2007)). It is interesting to note that word recall didn’t interfere with finger tapping in their study but it certainly did with an interval discrimination task in other experiment Miller, Hicks, and Willette (1978). This might suggest that we have to distinguish between different types of temporal tasks as it seems that they possibly don’t share the same cognitive resources. Finally, it is also possible that finger tapping could be performed with a high degree of automaticity, and therefore there would be resources left to perform a secondary task, such as the one used by Miyake *et. al*, without interfering with the main task. This hypothesis could be probed by increasing the difficulty of the secondary task.

We found evidence that tapping in NORMAL attention is done with a certain degree of automaticity, since the mental workload reported by the subjects is lower than in HIGH. In view of these findings, we can consider that finger tapping is not a strongly automatic task, in the sense that Kahneman and Chajczyk define it: a task whose performance doesn’t improve when attention is oriented towards it and doesn’t get worse when attention is diverted from it Kahneman and Chajczyk (1983). Instead, we can conclude that finger tapping fits better in the partially automatic category because of both its automaticity characteristics and its susceptibility to attention. According to them, a partially automatic task can be accomplished without attention although its performance improves when attention is oriented towards it.

The resynchronization phase is also affected by attention. We found that focusing the attention on the tempo change increases the resynchronization efficiency. This result is consistent with the study by Repp *et al*. Repp and Keller (2004), in which he observes a slower correction (smaller PCR) for conditions in which attention is diverted from the tempo change. According to the prior entry hypothesis, which states that an attended stimulus is detected faster than an unattended one Spence and Parise (2010), we can suppose that subjects in HIGH perceive the tempo change earlier than subjects in NORMAL, leaving them with more time for the processing. This may be one of the causes explaining the differences observed in resynchronization efficiency between the attention conditions.

### B. Neural correlates of attention in SMS

We found differences in the early components of the auditory-related ERPs between levels of attention. Such differences have been interpreted in the literature as an effect of attention Teder *et al*. (1993); Alho *et al*. (1994a); Hansen and Hillyard (1980); Eimer and Forster (2003a) and are compatible with the negative processing wave, which is a negative displacement of the attended-ERP versus the non-attended-ERP (HIGH and NORMAL in our work, respectively). The negative processing wave begins in the ascending deflection of N1; it may last a few hundred milliseconds and appears mainly in the front-central region Näätänen (1982). In an auditory and dichotic stimuli paradigm, Teder and coworkers report an amplitude modulation at P2 latencies in the same direction as ours: the amplitude of P2 in the attended-ERP is significantly lower than in the unattended-ERP (see Teder *et al*. (1993), Figures 1 y 2, ISI 480 ms). This gives support to our claim that in our task we are indeed controlling the level of attention. It is important to note that the stimuli sequence is the same for both levels of attention (as it only includes the pre-perturbation region), thus any differences in the ERPs may be attributed unequivocally to different psychological states.

### C. SMS and auditory feedback

Auditory feedback decreases the mean asynchrony in the pre-perturbation region, in line with Aschersleben’s report Aschersleben and Prinz (1995), although the effect is not significant. We suppose that this difference with Aschersleben’s results is due to the type of phones we used. Half of the subjects (no EEG recording) wore headphones which suppressed most of the surrounding noise; the other half (EEG recording) used earphones because the EEG didn’t leave enough space for headphones. Although a statistical test revealed no significant effect of phone-factor on asynchrony (see Methods), the earphone group was only partially isolated from the surrounding noise and the direct sound of taps was more audible, a situation that we believe contributed to diluting the effect of the auditory feedback when it was added.

We found that the resynchronization efficiency decreases when auditory feedback is added. From a naive point of view, this result is unexpected since the auditory feedback would give the subject access to more or better information. We hypothesize that stimuli and feedback are interfering with each other as both are in the same sensory modality and temporally overlapped. It is possible that auditory feedback prevents the subject from recognizing promptly the new stimuli period, and as a consequence he/she might be unable to perform the corrections efficiently.

### D. Asymmetric behavior in SMS

We observed that positive perturbations have a higher resynchronization efficiency than negative perturbations of the same magnitude. This asymmetry is in accordance with results from different paradigms (e.g. perception vs production), different perturbation types, and different measures to evaluate the resynchronization process Repp (2001, 2002b,a); Repp and Keller (2004); Repp (2011); Praamstra *et al*. (2003); Bavassi, Tagliazucchi, and Laje (2013); Jang *et al*. (2016). This asymmetry could be a signature of a non-linear correction mechanism Bavassi, Tagliazucchi, and Laje (2013); López and Laje (2019). Jang *et al*. poses an interesting hypothesis for this asymmetry Jang *et al*. (2016); but there is no agreed explanation for positive perturbations having higher resynchronization efficiency.

Our results for ∆*MA* (Figure 4A) show that a step-change period perturbation has its own effect on mean asynchrony, and it is opposite to the effect of parametrically varying a constant period. The effect on average is greater for negative perturbations; all in agreement with previous results Bavassi, Tagliazucchi, and Laje (2013). Our main conclusion from this result is that it reveals that post-perturbation MA is a variable to be controlled since it depends on the three factors: attention, feedback, and perturbation sign. Therefore, if the resynchronization process is to be evaluated it is important to define a magnitude that could comprise this effect. This is the reason that encouraged us to define the resynchronization efficiency, a magnitude containing in its definition the change in post-perturbation MA. Finally, the fact that a perturbation to the period has its own effect on the MA speaks against the inclusion of more than a single perturbation per trial, a common practice in some experimental designs (e.g. Jang *et al*. (2016); Praamstra *et al*. (2003)).

## V. AUTHOR CONTRIBUTIONS

LV and RL conceived the study; LV and RL designed the experiments; LV conducted the experiments and performed data analysis; all authors interpreted the data and wrote the manuscript.

## VII. ACKNOWLEDGMENTS

We thank Gustavo Juantorena and Pablo Riera for technical assistance and advice on the EEG recordings. We thank Claudia R. González for helpful comments on a previous version of the manuscript. We thank Rosanna Nicoletti for language editing.

## IX. DATA AND CODE AVAILABILITY

Data and code are available at the Sensorimotor Dynamics Lab website: http://ldsm.web.unq.edu.ar/attention/.

## Appendix A TapAdjar: A algorithm to remove distortion due to temporally adjacent ERPs

### 1. Introduction

In paced finger tapping with stimuli periods of a few hundreds milliseconds, the subject probably monitors his/her own performance from the last tap while engaging in motor preparation/execution for the next tap and tracking any change in the stimuli sequence López and Laje (2019); Bavassi *et al*. (2017); Bavassi, Tagliazucchi, and Laje (2013). The neural processes involved have durations comparable to the stimuli period, so some degree of overlapping between the response- and stimulus-related ERPs is expected. It would be desirable to have a method to separate both contributions in order to obtain an estimation of the isolated stimuli-related ERPs. By attenuating the distortions caused by the naturally overlapping response-related component, the isolated ERPs could be compared with the extensive literature on attention and auditory stimuli Näätänen (1982). To this end, in this appendix we propose an adaptation of an existing algorithm used in other experimental paradigms.

We base our proposal on Adjar, an algorithm designed by Woldorff Woldorff (1993) for a selective attention paradigm with auditory and dichotic stimuli in conditions with short ISIs of 300 ms and shorter. Considering such values of ISIs, an overlap between the ERPs generated by the present and the subsequent stimuli is expected to occur. This algorithm is meant to attenuate the distortions due to adjacent ERPs, allowing in this way an estimation of the ERP that could be obtained if there were no overlap.

We now describe our adaptation of Adjar to paced finger tapping; we call it *TapAdjar*. It is based on the idealization that each epoch recorded by the EEG primarily consists of two large, overlapping components that cannot be measured directly: one is related to the stimulus (beep) and the other is related to the corresponding response (tap) (Figure 6A). The general idea is to estimate the “pure” response-related ERP (Figure 6B) and to subtract it from the recorded ERP (Figure 6C); the result will be a stimulus-related ERP estimate with attenuated contamination from response-related sensorimotor potentials.

**FIG. 6.**
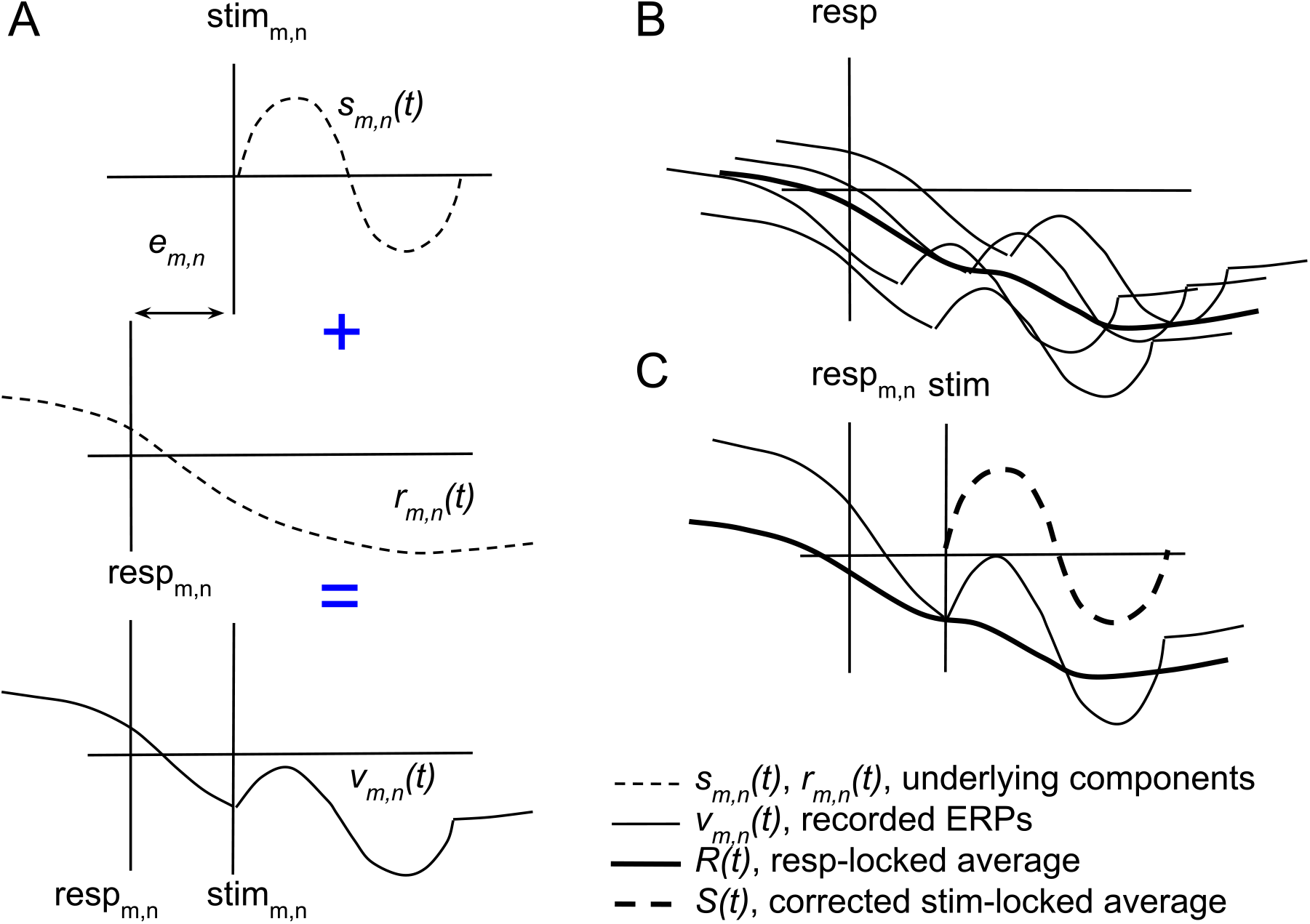
(A) Underlying (inaccessible) components in the *m, n*-th epoch (trial *m*, beep *n*). *s*_*m,n*_(*t*) is the auditory/stimuli-related component; high-frequency content. *r*_*m,n*_(*t*) is the response-related component; high- and low-frequency content. *v*_*m,n*_(*t*) = *s*_*m,n*_(*t*) + *r*_*m,n*_(*t – e*_*m,n*_) is the recorded epoch. *e*_*m,n*_ is the asynchrony of *m, n*-th epoch. (B) Single epochs *v*_*m,n*_(*t*) are aligned to taps and averaged, thus attenuating the high-frequency content. The result *R*(*t*) is the average response, that is an estimate of *r*_*m,n*_(*t*). (C) *R*(*t*) is shifted in time according to *e*_*m,n*_ and subtracted from *v*_*m,n*_(*t*) to obtain 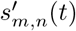 (not shown). This is repeated for all epochs and the results 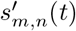 are aligned to the stimulus and averaged to obtain *S*(*t*) as the estimate of the auditory stimulus-related ERP.

### 2. Procedure

#### a. Step 1

We consider the pre-perturbation region only. We define the *m, n*-th epoch as the recorded EEG activity around the *n*-th stimulus (beep) and its corresponding response (tap) of the *m*-th trial, for each electrode, subject, and condition. The asynchrony corresponding to the *m, n*-th epoch is the temporal difference between the onset of the *m, n*-th response and the *m, n*-th stimulus, 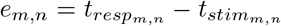. We assume that the recorded EEG is the superposition of two main components (Hypothesis I): a) a voltage variation related to both the *m, n*-th stimulus and the effects of attention; we call this component *s*_*m,n*_(*t*); b) a voltage variation related to the sensorimotor activity of the *m, n*-th tap; we call this component *r*_*m,n*_(*t*) (Figure 6A). These two components are usually overlapped, a condition that turns them individually inaccessible when using a single direct measure. Considering the *e*_*m,n*_ asynchrony between the onsets of response and stimulus, the EEG voltage of *m, n*-th epoch can be written as

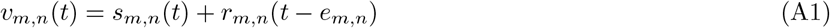

Then, the epochs *v*_*m,n*_(*t*) are aligned to responses and averaged:

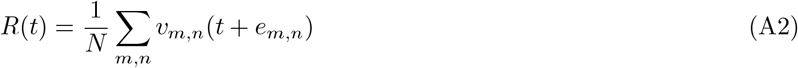

where *N* is the total number of epochs for each electrode, subject, and condition; *R*(*t*) is the commonly called averaged resp-locked ERP (Figure 6B). We consider *R*(*t*) to be an estimate of *r*_*m,n*_(*t*), the “pure” response-related component (Hypothesis II).

#### b. Step 2

We estimate *s*_*m,n*_(*t*), the component related to the auditory *m, n*-th stimulus. From equation A1 we obtain

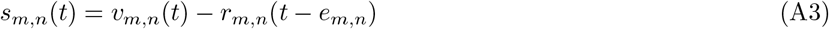

where *r*_*m,n*_(*t*) can be replaced by its estimation *R*(*t*) (Hypothesis III) appropriately shifted in time according to the asynchrony of that epoch. The result is 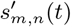, an estimation of *s*_*m,n*_(*t*):

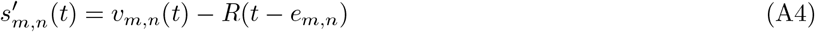

The process is repeated for each of the *N* epochs. Finally, the resulting 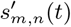 are aligned to the stimulus and averaged, obtaining the “corrected” averaged stim-locked ERP, or in other words, the estimate of the “pure” ERP related to the auditory stimuli (Figure 6C):

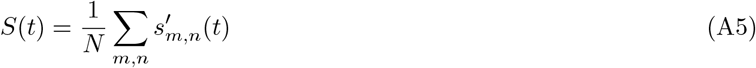

### 3. Justification of the hypotheses

Our proposal requires us to assume some hypotheses that, with the exception of the first one, are equivalent to those assumed by Woldorff. Here, we describe them using the terminology of finger tapping.

**Hypothesis I**. *Each epoch is composed of only two large components: one related to the stimulus and the other related to the corresponding response. We also consider that each epoch is the sum of the components pertaining only to that epoch, and we assume that there is a negligible overlap with adjacent epochs (first order of approximation, according to Woldorff terminology Woldorff (1993))*.

In a finger tapping task, there is an overlap between the EEG activity related to stimulus and response Praamstra *et al*. (2003). Despite the low-pass filtering effect of epoch averaging due to asynchrony variability, the stim-locked ERPs still show residuals resulting from the sensorimotor activity associated with the responses. This residual activity is clearly observed before the stimulus onset as a low-frequency negative deflection of the potential, and on later latencies it is found overlapped with the auditory ERP components Praamstra *et al*. (2003); Robinson and Rudge (1977). According to the literature, it is reasonable to suppose that this low frequency deflection is the post-movement potential observed by Gerloff (Gerloff *et al*. (1997), Figure 2).

In addition to the components related to finger movement, we have to consider the activity generated by other causes. For example, when the finger hits the sensor a tactile ERP is generated, presumably with frequencies and latencies similar to those of the auditory ERPs Eimer and Forster (2003a,b). Besides, in conditions with additional auditory feedback we should take into account a response-related auditory ERP. In summary, the post-movement potential ERP, the tactile ERP, and the auditory-feedback ERP can be considered related to the response.

As for the stimulus-related component, we take into account the auditory-stimulus ERP and the effect of attention. We could also consider the appearance of a readiness potential or even a contingent negative variation (CNV) relative to the sensor-switching movement of the finger (it takes place once per trial and only in HIGH conditions, see Methods), yet our results didn’t reveal any deflection compatible with those potentials. Finally, the sensor-switching movement occurs outside the region in which we are comparing ERPs (pre-perturbation region), so no potentials related to the sensor-switching movement are considered.

**Hypothesis II**. *When the number of averaged epochs is large enough, the resp-locked-ERP presents a negligible contamination from the stimulus-related components*.

As stated above, the stim-locked ERPs present a low-frequency negative deflection clearly visible previous to the onset of the stimulus. This deflection is a consequence of the components related to the tap Praamstra *et al*. (2003). We suppose that the activity related to the tap has both low-frequency components (due to the post-movement potential) and high-frequency components coming from the other contributions. Instead, the activity related to the stimuli has a diminished low-frequency content Praamstra *et al*. (2003). This asymmetry in the frequency content between stimuli and taps has an important consequence: when epochs are aligned to the responses and averaged, high frequencies coming from the stimuli are averaged out by the natural variability of asynchronies. This fact makes it possible to isolate or, at least, obtain an estimate of the “pure” resp-locked ERP. On the contrary, when epochs are aligned to stimuli and averaged, the asynchrony variability again smooths high frequencies in the tap-related activity, but not the low ones, which persist in the stim-locked ERP.

**Hypothesis III**. *In terms of EEG activity, we assume that all the responses for a given subject, condition, and electrode are equivalent; and that the averaged resp-locked ERP can be considered as an estimation of each individual response-related component; that is, R*(*t*) ∼ *r*_*m,n*_(*t*).

We will now consider three possible causes of differences in individual response-related activity: overlap of adjacent epochs, EEG noise, and asynchrony value; we now show reasons to safely ignore their effect. Regarding the first possible cause, an ISI as the one used in this work (600 ms) is considered large enough to minimize overlap between adjacent epochs Alho *et al*. (1994b); Teder *et al*. (1993); Woldorff (1993), so we only consider that there exists overlap between stimulus and tap activities belonging to the same epoch; possible overlaps between epochs are considered as higher order corrections and so we ignore them (Hypothesis I). The second reason that could cause two individual responses to be different is EEG noise, normally modeled as white noise Woldorff (1993); therefore we expect that when several epochs are averaged its random effect is considerably reduced. Finally, the naturally variable asynchrony value might make two individual responses different. That is, the EEG activity associated to a response whose asynchrony is –250 ms might be different to the one associated to a response whose asynchrony is +150 ms. Yet, as it was said in the main text, epochs with extreme values of asynchrony were discarded. We kept epochs with asynchronies between –170 and +80 ms, what reduces the possible variability of ERPs because of this reason.

### 4. Results

Figure 7 shows the grand-average ERPs for NORMAL and HIGH conditions (feedback conditions averaged) corresponding to the 8 electrodes in the front-central region (AF3, F3, F7, FC5, FC6, F8, F4, AF4). Panel B shows the stim-locked ERPs before any corrections were made. It can be observed a low-frequency negative deflection of the potential previous to the onset of the stimulus (dashed vertical line). Praamstra gave evidence that this deflection is caused by the activity related to the tap Praamstra *et al*. (2003). In later latencies, the superposition of the activity coming from the tap makes the stim-locked ERPs appear distorted with respect to the auditory canonical ERP shape, with the three well defined peaks, P1, N1, and P2. Panel A shows the resp-locked ERPs. A bootstrapped comparison shows no significant differences between NORMAL and HIGH ERPs in a window going from –100 to +300 ms relative to the onset of the tap. Panel C shows the stim-locked ERPs after the TapAdjar correction was applied (as plotted in Figure 5A). The TapAdjar algorithm was applied to each subject, condition, and electrode before the average process was performed. We can observe now that the region previous to the onset of the stimulus doesn’t show the negative deflection anymore, and although the ERPs don’t show a flat baseline, at least both ERPs appear to rise from near zero voltage values, which is convenient for comparing absolute amplitude values between conditions. Also, we can observe a P1 peak which is not present in the stim-locked ERPs before correction, a fact that increases the similarity of the corrected ERPs with those elicited by an isolated auditory stimulus. Error bands in corrected ERPs are narrower than in uncorrected ERPs; this suggests that the TapAdjar algorithm doesn’t have a random effect over the ERPs.

**FIG. 7.**
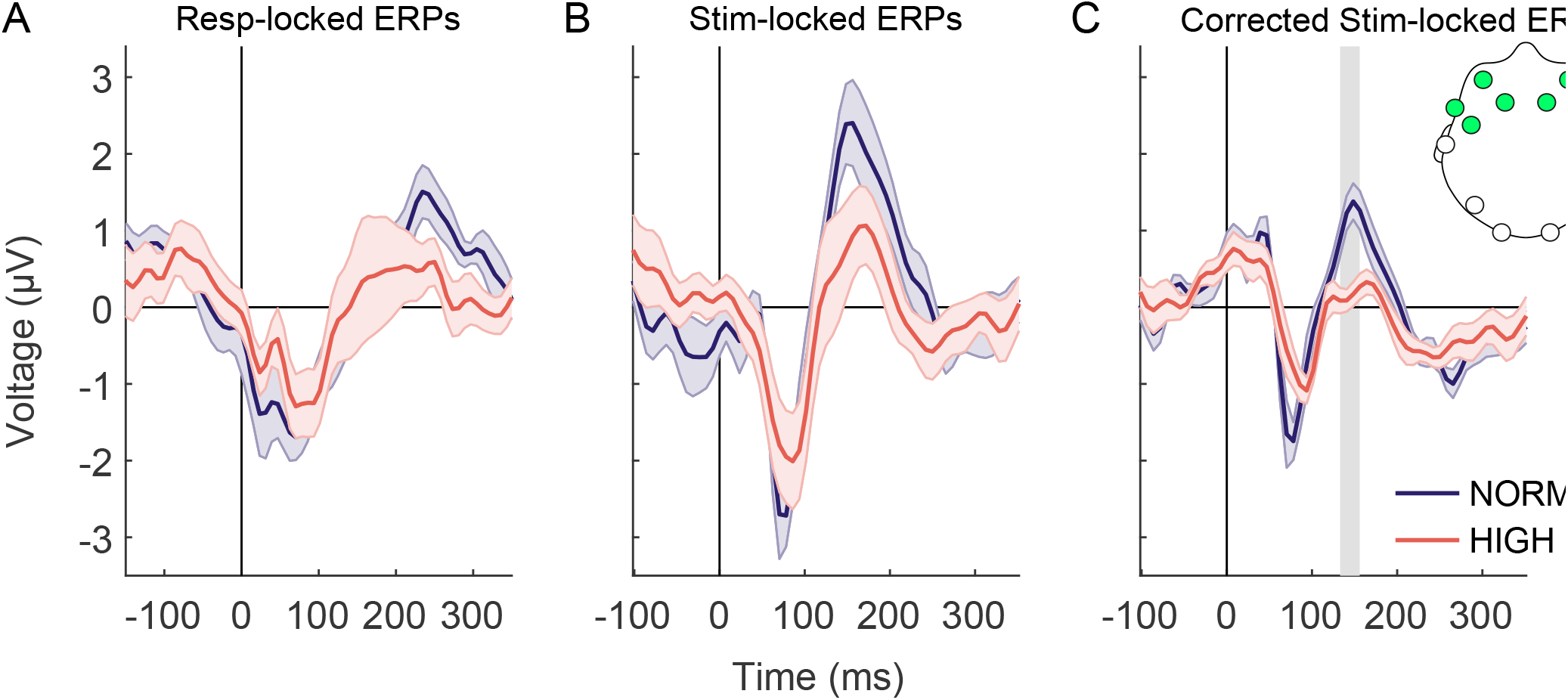
(A) Recorded resp-locked ERPs. (B) Recorded stim-locked ERPs. A low-frequency negative deflection is observed near the onset of the stimulus. (C) Corrected stim-locked ERPs (as plotted in Figure 5A). The negative deflection now is absent, and while the baseline is not flat at least both ERPs begin to rise from near zero voltage values. Also a first peak associable with P1 is observed, as it is expected from a canonic auditory ERP. **Notes:** In panel A, the vertical dashed line represents the onset of the tap; in panels B and C, represents the onset of the stimulus. The three panels show the general mean ERPs, where feedback conditions were averaged. The TapAdjar correction was applied for each subject, condition, and electrode, before the averaging process was performed. Mean *±* standard error across subjects.

